# Transcriptional Profiling of Commonly Used Liver Cancer Cell Lines Reveals Disease-Specific Modeling Potential and Authentication Concerns

**DOI:** 10.1101/2025.10.30.685599

**Authors:** Nima Sarfaraz, Gail Rosen, Trevor Winkler, Risha Patel, Srinivas Somarowthu, Michael Bouchard

## Abstract

Cell lines are essential tools for liver cancer research, yet their molecular fidelity to primary tumors remains incompletely characterized. Here we comprehensively evaluated transcriptomic similarities between commonly used liver cancer cell lines and primary tumor subtypes to guide optimal model selection. We analyzed RNA sequencing data from 541 samples spanning primary HCC, HPBL, CHOL, and FLC tumors, alongside 21 liver cancer cell lines and primary human hepatocytes. Through systematic variance analysis, we identified 2,523 highly variable genes distinguishing cancer subtypes and cell lines, then performed correlation analyses, unsupervised clustering, and pathway enrichment to assess molecular similarities. Molecular subtypes within each cancer type were identified through hierarchical clustering and characterized using pathway analysis. HepG2 cells showed strongest correlation with HPBL (r=0.62), confirming their hepatoblastoma origin despite frequent HCC misclassification. This correlation was driven by shared Wnt pathway dysregulation signatures. Huh7 cells best represented HCC, particularly the immune-modulatory, MYC-activated subtype with the highest median correlation. RBE cells optimally modeled CHOL, specifically the dedifferentiated, immune-evasive subtype. Several commonly used cell lines (LO2, SMMC-7721, MHCC97L) and specific publicly available samples demonstrated likely HeLa contamination. Primary human hepatocytes cultured under physioxic conditions better preserved liver-specific transcriptional programs compared to standard culture. No established cell line analyzed represented FLC strongly, identifying the need for a standard, available model. This transcriptomic framework provides evidence-based guidance for selecting appropriate liver cancer cell line models and highlights the critical need for rigorous cell line validation to improve experimental design and translational relevance of liver cancer research.

## Introduction

Primary liver cancer represents a significant global health challenge, ranking as the third leading cause of cancer-related deaths worldwide.^1^ This disease encompasses several distinct subtypes, each characterized by unique molecular features and clinical outcomes. Hepatocellular carcinoma (HCC), accounting for approximately 85% of cases, typically develops in the context of chronic liver disease.^1^ Intrahepatic cholangiocarcinoma (CHOL), originating from bile duct epithelium, is the second most common subtype. Less frequent variants include fibrolamellar carcinoma (FLC) and hepatoblastoma (HPBL), which predominantly affect younger populations.^2,3^ The molecular and clinical heterogeneity across these subtypes highlights the complexity of liver cancer and the need for tailored approaches in both research and treatment strategies. Within each liver cancer type exists substantial heterogeneity to consider as well, presenting additional challenges for developing representative experimental models and emphasizes the importance of careful model selection for specific research questions.^4,5^

In vitro cell lines have long served as invaluable tools for studying the molecular mechanisms underlying liver cancer and for developing potential therapeutic strategies. These models offer researchers a controlled environment to investigate cancer biology, drug responses, and cellular processes, providing insights that can ideally be translated into in vivo and clinical settings. Given the complexity and diversity of primary liver cancers, it is crucial to select in vitro models judiciously and understand to what degree they represent and recapitulate the biological characteristics of the disease systems of interest. As such, the choice of cell line model should be guided by the specific research context and molecular characteristics required, as different experimental objectives demand careful consideration of distinct cellular properties.

Context-dependent model selection is critical in liver cancer research, exemplified across multiple key applications. Huh7 cells are the primary HCV model due to exceptional viral replication support, though this varies between subclones through differential expression of host factors like THAP7 and NR0B2.^6^ Hep3B and PLC/PRF/5 harbor integrated HBV DNA and produce viral surface antigens, making them valuable for studying HBV-associated HCC and HDV infection.^7,8^ However, their endogenous HBx expression modulates NF-κB signaling, p53-mediated apoptosis, and DNA damage responses, requiring careful consideration in pathway analyses.^9^ Drug metabolism studies demand evaluation of metabolic competencies across models. While HepG2 and HepaRG cells express phase I and II drug-metabolizing enzymes, both show reduced capacity compared to primary hepatocytes, particularly for specific CYP isoforms.^10,11^ Cell line selection also impacts signal transduction studies due to baseline pathway differences. HepG2 cells display constitutive PI3K/Akt activation, while Huh7 cells exhibit elevated MAPK signaling.^12–15^ These inherent differences, combined with cell line-specific mutations like p53 alterations in Hep3B, can significantly affect interpretation of pathway perturbation experiments and therapeutic responses.^16^

Careful interpretation of cell line models can be compromised by misclassifications and misuse in the literature, introducing confounding factors and highlighting the need for rigorous validation. A prominent example is HepG2 cells; despite numerous publications and the ATCC citing HepG2 as HCC-derived, investigations revealed the original source tissue was clinically classified as HPBL.^17^ This misattribution has created a self-reinforcing cycle, with subsequent studies perpetuating the incorrect HepG2-HCC classification. Consequently, HepG2 cells have been used to model HCC-specific outcomes, potentially causing misinterpretation in adult liver cancer biology contexts, while being overlooked for HPBL-relevant studies. Further complicating the landscape of liver cancer cell models are instances of cell line contamination or misidentification. LO-2, originally described as fetal hepatocyte-derived, and SMMC-7721, supposedly HCC-derived, have been reported as cervical adenocarcinoma HeLa cells or HeLa-contaminated.^18–20^ These findings reinforce the critical importance of cell line authentication and the far-reaching implications of using misidentified or mischaracterized cell lines in research.

Towards this effort, numerous studies have sought to profile and characterize these cell line models to provide information beyond their original clinical annotations. These efforts have employed various approaches, including proteomic analyses to elucidate protein expression patterns and post-translational modifications, DNA sequencing to identify chromosomal aberrations and driver mutations, STR profiling, and phenotypic assays to assess and compare functional characteristics such as proliferation rates, invasion capacity, and drug sensitivity.^10,21–23^ While these studies provide valuable information and insights, they have typically been limited in scope, focusing on a narrow range of endpoints and a limited number of cell lines, and often without directly comparing back to primary liver cancer subtypes.

Recent high-throughput RNA-sequencing advances have enabled comprehensive transcriptomic analyses of liver cancer cell lines. Studies comparing cell lines like HepG2 and HepaRG with primary liver tissue have revealed significant differences in gene expression profiles and pathway coverage, emphasizing the importance of careful model selection.^10^ Comparative analyses of hepatoma cell lines such as HepG2 and Huh7 against primary human hepatoma cells and hepatocytes have uncovered important distinctions in gene expression patterns and metabolic characteristics, demonstrating that cell lines may not always accurately reflect primary liver cancer molecular features.^6,21,24^ Studies comparing HepG2 and Hep3B have revealed significant differences in gene expression, drug responses, and signaling pathways despite their frequent interchangeable use.^22^ These findings reinforce the importance of understanding the unique characteristics of each cell line and their potential limitations in modeling specific aspects of liver cancer biology.

Despite these advances, there still remains a gap in comprehensive transcriptome-wide studies that directly compare liver cancer cell lines to the diverse subtypes of primary liver cancers and to each other. A more extensive analysis that integrates transcriptomic data from a broader range of cell lines with profiles from various liver cancer subtypes could provide crucial insights into the strengths and limitations of these models in representing specific disease states. Such comprehensive comparisons would not only enhance our understanding of how accurately these cell lines model different aspects of liver cancer biology but also guide researchers in selecting the most appropriate models for specific research questions or drug discovery efforts. Likewise, this approach could potentially identify novel cell line use cases that recapitulate the molecular features of liver cancer subtypes, ultimately leading to more translatable preclinical research in hepatocellular carcinoma and related liver malignancies.

Here, we utilize publicly available bulk tissue RNA-seq samples and present a methodology to characterize a wide array of commonly used liver cancer cell lines in relation to primary liver cancer subtype clinical samples and to each other. Through this transcriptome-centric approach, we aim to enhance the understanding of how liver cancer cell lines relate to their tumors of origin and to other cell lines. Ultimately, this study aims to serve as a contributing piece alongside other information as part of comprehensive material to support more informed use of cell lines in liver cancer research.

## Material and methods

### Sample Collection and Data Sources

Publicly available RNA sequencing data were obtained from the Gene Expression Omnibus (GEO) database to enable comprehensive analysis of liver cancer cell lines and primary liver cancer samples. The dataset comprised a total of 541 samples as detailed in the metadata in the supplemental files. Samples were selected based on stringent criteria to maintain data quality and comparability: a minimum sequencing depth of 20 million reads per sample, libraries generated from total RNA extractions, and only control or vehicle-treated samples (H_₂_O, DMSO) from experimental studies, along with baseline conditions from genetic perturbation experiments (siCtrl, empty vector controls), were included.

The dataset included primary patient tissue samples described as tumor biopsies from hepatocellular carcinoma (HCC), hepatoblastoma (HPBL), fibrolamellar carcinoma (FLC), cholangiocarcinoma (CHOL), disease-free liver tissue (Liver), cervical squamous cell carcinoma (CESC), and normal cervical tissue (Cervix). Cell lines included in the analysis were HepG2, Hep3B, Huh7, Huh6, RBE, SNU475, SNU1079, TFK1, SKHep1, PLC/PRF/5, Primary Human Hepatocytes (PHHs), HeLa, SMMC-7721, MHCC97H, MHCC97L, and LO-2. For PHH samples, these were further categorized based on culture conditions: fresh (uncultured), short-term (maximum one passage and up to 48h in culture), short-term physioxic (maximum one passage at 7% O_₂_ and up to 48h in culture), and long-term (five passages and up to 45 days in culture). The inclusion of cervical cancer samples and HeLa cells served dual purposes: they provided external reference points for validating the clustering methodology and enabled the detection of potential HeLa cell contamination previously reported in some liver cancer cell lines.

### RNA Sequencing Data Processing

Raw sequencing files (FASTQ format) were downloaded using SRA Toolkit v3.0.0. Quality control of raw reads was performed using FastQC (v0.11.9), and MultiQC (v1.11) was used to aggregate quality reports. RNA sequencing data were processed using the Nextflow nf-core/rnaseq pipeline, STAR for read alignment to the human reference genome (GRCh38), and StringTie for transcript assembly and quantification.

Gene-level count matrices were generated using StringTie’s prepDE.py script, resulting in raw count matrices suitable for differential expression analysis. The DESeq2 package (v1.34.0) in R (v4.2.0) was used to create a DESeqDataSet object from the count matrix. Variance stabilizing transformation (VST) was applied to normalize the count data for downstream analyses.

To mitigate potential batch effects arising from different experimental sources, we employed ComBat-Seq from the sva package (v3.42.0) on the raw counts matrix prior to DESeq2 normalization. This approach was applied separately to each tissue/cell line group containing samples from multiple bioprojects, using bioproject as the batch variable while preserving biological variation associated with tissue identity. The effectiveness of batch correction was assessed using Principal Component Analysis (PCA) on the variance-stabilized transformed data. Visual inspection of PCA plots before and after correction confirmed the reduction of bioproject-associated clustering within each tissue type, while maintaining separation between biologically distinct groups. For tissue/cell line groups with only one sample per bioproject, the original counts were retained to avoid introducing artifacts.

### Variance Analysis and Marker Gene Identification

To identify genes most informative for comparative analyses between cell lines and primary tumors, we first addressed a fundamental challenge in cross-sample transcriptome correlative comparisons: when analyzing only using the full gene set, samples typically show high cross-correlations regardless of etiology, primarily reflecting shared basic human cellular processes and origin rather than biologically meaningful differences. This high baseline similarity can mask subtle but important biological distinctions, paralleling challenges in other genomic analyses, where focusing on informative features - such as variants in population genetics ^26^, marker genes in single-cell transcriptomics ^27^, or selected features in machine learning approaches - enables detection of meaningful biological differences that would otherwise be obscured by shared genomic backgrounds. To increase sensitivity for detecting context-specific differences, we implemented a systematic approach to identify marker genes showing high biological variation while controlling for technical aspects of RNA-seq data.

We first filtered genes for reliable detection in cell line models by selecting those with mean expression above the 25th percentile. This initial filtering step ensured downstream analyses would capture meaningful biological variation while minimizing technical noise from lowly-expressed transcripts. We then conducted parallel analyses of expression variability across defined groups of cell lines and primary tumors:

For cell lines, we selected a balanced panel representing distinct reported disease origins, including HCC (Hep3B, Huh7, MHCC97H, SNU475, PLC/PRF/5), cholangiocarcinoma (RBE, SNU1079, TFK1), hepatoblastoma (HepG2, Huh6), and other cancer types (SK-Hep1, HeLa). For primary tumors, we focused on primary liver cancer subtypes as the purpose of this study was to anchor around their distinguishing characteristics.

For each gene, we calculated the mean expression and coefficient of variation squared (CV²) using group-level average expression profiles, treating each disease subtype or cell line category as a distinct group. The relationship between mean expression and variation was visualized using scatter plots. To model expected variance levels for a given mean expression, we fitted a robust linear regression with a 95% confidence interval. Genes showing higher-than-expected variability across groups were identified based on their deviation from the fitted regression model. Specifically, genes falling above the 95% confidence interval were considered to have significant biological variation, assessed using chi-square tests and false discovery rate (FDR) correction (adjusted p-value < 0.05).

This analysis identified 11,052 highly variable genes among cell lines and 4,395 among primary tumors. The intersection of these sets yielded 2,523 genes showing significant variation across unique groups in both contexts. This shared signature of highly variable genes formed the basis for subsequent analyses comparing expression patterns between liver cancer cell lines and their primary tumor counterparts.

### Pairwise Correlations

Pairwise Spearman correlations were calculated between averaged gene expression profiles of each tumor type and cell line, using the identified marker genes. For more detailed analysis, individual correlations were computed between each cell line sample and each primary tumor sample. Correlation matrices were generated to visualize the similarities among all sample groups. For statistical robustness, median correlation values were calculated across all pairwise sample comparisons within relevant groups. To evaluate individual cell line samples, we developed a violin plot approach that visualized the distribution of correlations between all samples of a given cell line and all samples of each tumor type. This approach enabled identification of potential sample heterogeneity or outlier contamination, which was subsequently confirmed through additional analyses.

### Hierarchical Clustering

Unsupervised hierarchical clustering was performed using the marker gene set to explore relationships among samples. Euclidean distance was calculated on variance-stabilized expression values as the distance metric, and Ward’s minimum variance method (ward.D2) was employed as the linkage criterion to minimize within-cluster variance. Clustering was visualized using pheatmap (v1.0.12), with sample groups annotated by tissue type and origin. For gene clustering, the same distance metrics and linkage methods were applied to identify co-expressed gene modules.

### Principal Component Analysis

PCA was performed on VST-normalized expression data of the marker gene set to visualize sample relationships in a reduced-dimensional space. PCA was implemented using the plotPCA function in DESeq2, with the top 500 most variable genes used to calculate the principal components. The first two principal components, which captured the greatest proportion of variance, were plotted to visualize the clustering patterns of samples. Probability ellipses were calculated and overlaid to delineate the 95% confidence regions for each sample group, providing a statistical assessment of group separation.

### Sample Authentication and Contamination Detection

Correlation patterns of individual samples were assessed to identify potential contamination or misidentification issues. Samples showing unexpected correlation patterns were flagged and segregated into anomalous categories (e.g., Hep3B_Anom, Huh7_Anom) for separate analysis.

### Molecular Subtype Identification

To identify molecular subtypes within each liver cancer type and assess how well cell lines model these distinct tumor phenotypes, we employed a multi-step analytical approach:

For each cancer type (HCC, HPBL, CHOL, and FLC), we performed unsupervised clustering using highly variable genes identified through variance modeling of normalized expression data across all samples of that cancer type. Gap statistic analysis was used to determine the optimal number of clusters, with bootstrap resampling (n=50) to ensure statistical robustness.

Hierarchical clustering with Ward’s linkage method was applied to identify discrete molecular subtypes within each cancer type. Cluster stability was assessed using silhouette analysis to ensure robust subtype identification. The biological characteristics of each identified subtype were determined through differential expression analysis comparing samples within a cluster to all other samples of that cancer type.

To characterize the molecular features of each subtype, we employed multiple pathway analyses. Gene Ontology (GO) enrichment identified enriched biological processes in cluster-specific gene signatures. KEGG pathway analysis revealed altered signaling networks characteristic of each subtype. Gene Set Enrichment Analysis (GSEA) determined activation or suppression of hallmark pathways relative to other clusters, providing insight into the functional state of each subtype.

To evaluate how well cell lines model specific tumor subtypes, we calculated correlation coefficients between each cell line’s gene expression centroid and the centroids of identified tumor clusters. These correlations were visualized through spider plot analysis, enabling identification of which tumor subtypes are best represented by each cell line model.

### Gene Set Enrichment and Pathway Analysis

Functional characterization of the marker gene set and subtype-specific genes was performed using multiple complementary approaches:

Gene Ontology (GO) enrichment analysis was conducted using the clusterProfiler package (v4.0.5) with the org.Hs.eg.db annotation database. GO terms were categorized into biological process (BP), molecular function (MF), and cellular component (CC) domains. Enrichment was assessed using a hypergeometric test with Benjamini-Hochberg correction for multiple testing (adjusted p-value < 0.05).

KEGG pathway analysis was performed to identify enriched biological pathways among the marker genes and subtype-specific genes. Enrichment was calculated using the enrichKEGG function from clusterProfiler with an adjusted p-value cutoff of 0.05.

### Pathway Level Analysis of Gene Expression (PLAGE)

To evaluate the preservation of key biological pathways across cell lines and tumor types, we employed Pathway Level Analysis of Gene Expression (PLAGE) using the GSVA package (v1.42.0). Selected gene signatures representing liver-specific functions (HSIAO_LIVER_SPECIFIC_GENES), drug metabolism (KEGG_DRUG_METABOLISM_CYTOCHROME_P450), metastatic potential (ROESSLER_LIVER_CANCER_METASTASIS_UP), and apoptotic processes (GOBP_HEPATOCYTE_APOPTOTIC_PROCESS) were analyzed across all samples. PLAGE scores were calculated for each signature and sample, then averaged across sample groups to identify patterns of pathway activity.

### Differential Pathway Activity in Molecular Subtypes

For each identified molecular subtype, we performed GSEA to determine differentially active pathways relative to other subtypes of the same cancer type. Gene lists were ranked by differential expression (using log2 fold change), and enrichment was calculated against the MSigDB Hallmark gene sets, C2 curated gene sets, and C5 GO gene sets. Normalized enrichment scores (NES) and adjusted p-values were calculated to identify significantly enriched pathways (adjusted p-value < 0.05).

## Results

### Sample Selection and Data Preparation

We analyzed 541 RNA-seq samples, as described in the methods, comprising primary liver tumors (HCC, HPBL, FLC, CHOL), normal liver tissue, cervical cancer controls (CESC, Cervix), and 21 liver cancer cell lines including primary human hepatocytes under various culture conditions (Fig 1, Supplemental Data). Cervical and HeLa samples served as external validation controls and enabled detection of potential HeLa contamination previously reported in liver cancer cell lines.^18,19,25^

**Figure 1.**
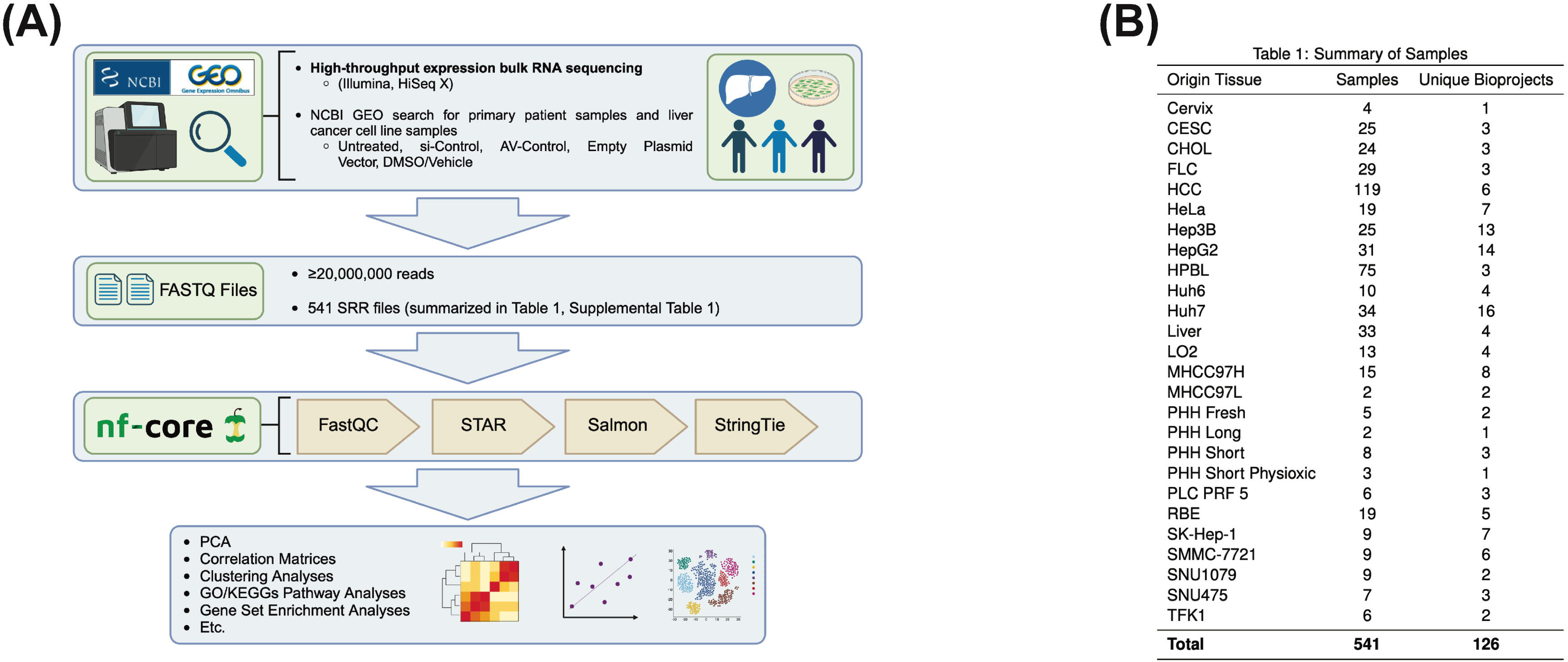
Study design and sample composition. This workflow integrates multiple public datasets to create a robust framework for evaluating how well commonly used liver cancer cell lines recapitulate the transcriptional features of primary tumors. (A) Workflow schematic for sample selection and processing. (B) Summary of sample groups included in the analysis.

### Identification and Characterization of Relevant Gene Marker Set

To identify genes distinguishing between liver cancer subtypes and cell lines, we implemented variance modeling on group-level expression profiles as described in the methods. After filtering for reliable detection (>25th percentile), we analyzed expression variability across cell line groups (HCC: Hep3B, Huh7, MHCC97H, SNU475, PLC/PRF/5; CHOL: RBE, SNU1079, TFK1; HPBL: HepG2, Huh6; Other: SK-Hep1, HeLa) and primary tumor types. Genes showing higher-than-expected variability (above 95% CI of fitted regression; chi-square test, FDR < 0.05) were identified as highly variable (Fig 2A). This yielded 11,052 variable genes among cell lines and 4,395 among primary tumors, with 2,523 genes overlapping (Fig 2B).

**Figure 2.**
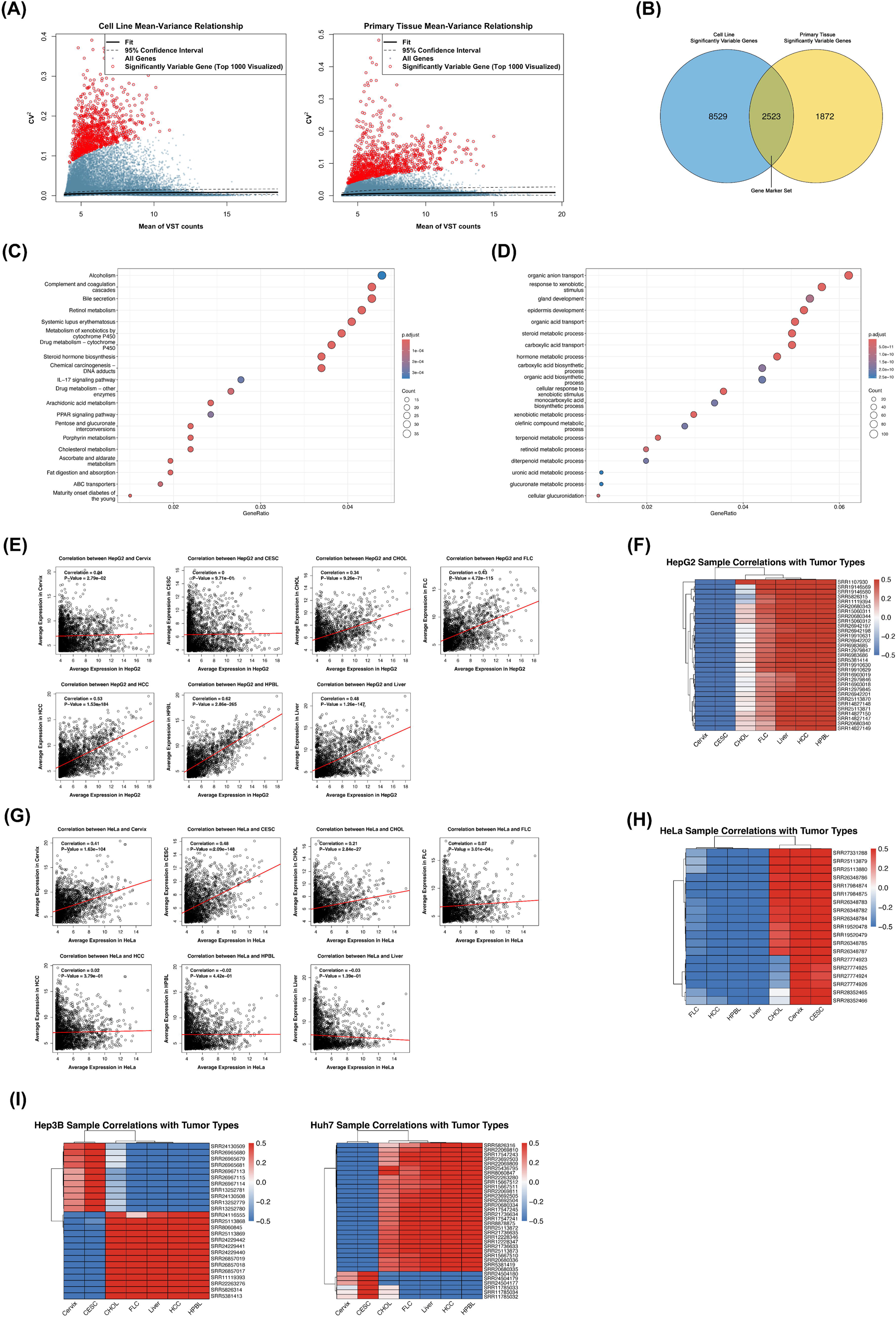
Identification and validation of a liver cancer transcriptional marker gene set. (A) Mean-variance relationship in VST-transformed expression data with fitted regression line (black) and chi-square confidence intervals (black dashed). Red points indicate genes with significant biological variation. (B) Venn diagram showing overlap between highly variable genes in cell lines and primary tumors. (C) Top enriched KEGG pathways in the marker gene set. (D) Top enriched Gene Ontology biological processes in the marker gene set. (E) Spearman correlation pattern of HepG2 cells across averaged gene expression profiles of tissue types using marker genes. (F) Individual HepG2 sample correlations to tissue types using marker genes, displayed as z-score scaled correlations in a heatmap. (G) Spearman correlation pattern of HeLa cells across averaged gene expression profiles of tissue types using marker genes. (H) Individual HeLa sample correlations to tissue types using marker genes, displayed as z-score scaled correlations in a heatmap. (I) Z-score scaled correlation heatmaps revealing HeLa-like samples within Hep3B (left) and Huh7 (right) collections, with clustering dendrograms indicating distinct groups of samples with HeLa-like transcriptional profiles. While most cell lines show expected correlation patterns with their reported tissues of origin, the identification of HeLa-like transcriptional profiles in subsets of Hep3B and Huh7 samples highlights the critical importance of molecular authentication in cell line research.

Pathway enrichment revealed these marker genes captured relevant liver-specific biology and cancer-related processes distinguishing tumor subtypes. KEGG analysis showed enrichment in metabolic pathways including alcoholism, complement and coagulation cascades, bile secretion, drug metabolism via cytochrome P450, and xenobiotic processing, reflecting both hepatic function and differential dysregulation across cancer types (Fig 2C).^28–31^ Gene Ontology analysis highlighted transport processes (organic anion/acid transport), developmental processes (gland/epidermis development), and metabolic pathways (steroid, retinoid, terpenoid metabolism) (Fig 2D). These pathways, essential for cellular homeostasis, show distinct patterns of dysregulation across different liver cancer subtypes.^32–37^ Collectively, this pathway analysis demonstrates that our marker gene set not only captures tissue-specific liver functions and cancer-related processes but also highlights the biological pathways that drive molecular differences between liver cancer subtypes and cell lines, validating their use as discriminatory markers for comparative analyses.

To validate our approach, we generated average expression profiles for each sample group and computed pairwise Spearman correlations. This analysis revealed correlation patterns largely consistent with known tissue origins. HepG2 cells showed strong-to-moderate positive correlations with liver-derived tissues, highest with HPBL samples (r = 0.62), while displaying weak or no correlations with cervical tissues and CESC (Fig 2E). Conversely, HeLa cells exhibited expected patterns for cervical cancer, with moderate-strong positive correlations to cervical and CESC tissues (r = 0.48) and weak or negative correlations to liver-derived samples (Fig 2G). To assess potential heterogeneity within cell line groups, we examined correlations at the individual sample level. HepG2 samples revealed consistent patterns across files, with all samples showing strong positive correlations to liver-derived tissues, particularly HPBL, and negative correlations to cervical tissues (Fig 2F). Similarly, HeLa samples demonstrated uniformly strong correlations to CESC and cervical tissues while displaying negative correlations to liver-derived samples (Fig 2H). While most individual samples met basic expectations, this granular approach revealed unexpected patterns within the Hep3B and Huh7 sample sets (Fig 2I, left and right respectively). Although most samples displayed expected correlation patterns, small subsets from both cell lines showed markedly divergent profiles. Several Hep3B samples (SRR24130509, SRR26965680, SRR26965679, SRR26965681, SRR26967113, SRR26967115, SRR26967114, SRR13252781, SRR24130508, SRR13252779, SRR13252780) showed correlation patterns highly similar to HeLa cells, diverging from the liver-specific signatures observed in their cohort. Similarly, a subset of Huh7 samples (SRR24504180, SRR24504179, SRR24504177, SRR11785033, SRR11785034, SRR11785032) displayed correlation patterns more characteristic of CESC tissue, deviating from the expected hepatocellular profile. These findings suggest potential HeLa cell contamination or misidentification within these sample sets, a known issue in cell culture work. ^25,39,40^ To maintain analytical rigor, we segregated these anomalous samples into distinct categories (Hep3B_Anom and Huh7_Anom). This reclassification preserved the integrity of the Hep3B and Huh7 expression profiles for our subsequent analyses while allowing us to separately explore the anomalous samples.

### Comprehensive Correlation and Clustering Analyses

An overarching PCA of individual samples using the signature gene set, along with a correlation matrix analysis of averaged group gene expression profiles (summarizing all comparisons as demonstrated in Figs 2E and 2G), revealed clear segregation of samples based on both tissue origin and disease state (Figs 3A and 3B). We further explored these relationships through pairwise Spearman correlations between all individual samples from each primary tumor group against all individual cell line samples ranked by median (Fig 3C) and an unsupervised hierarchical clustering based on the Euclidean distance matrix of averaged gene expression profiles per group (Fig 4A), to reinforce and expand upon observed patterns.

**Figure 3.**
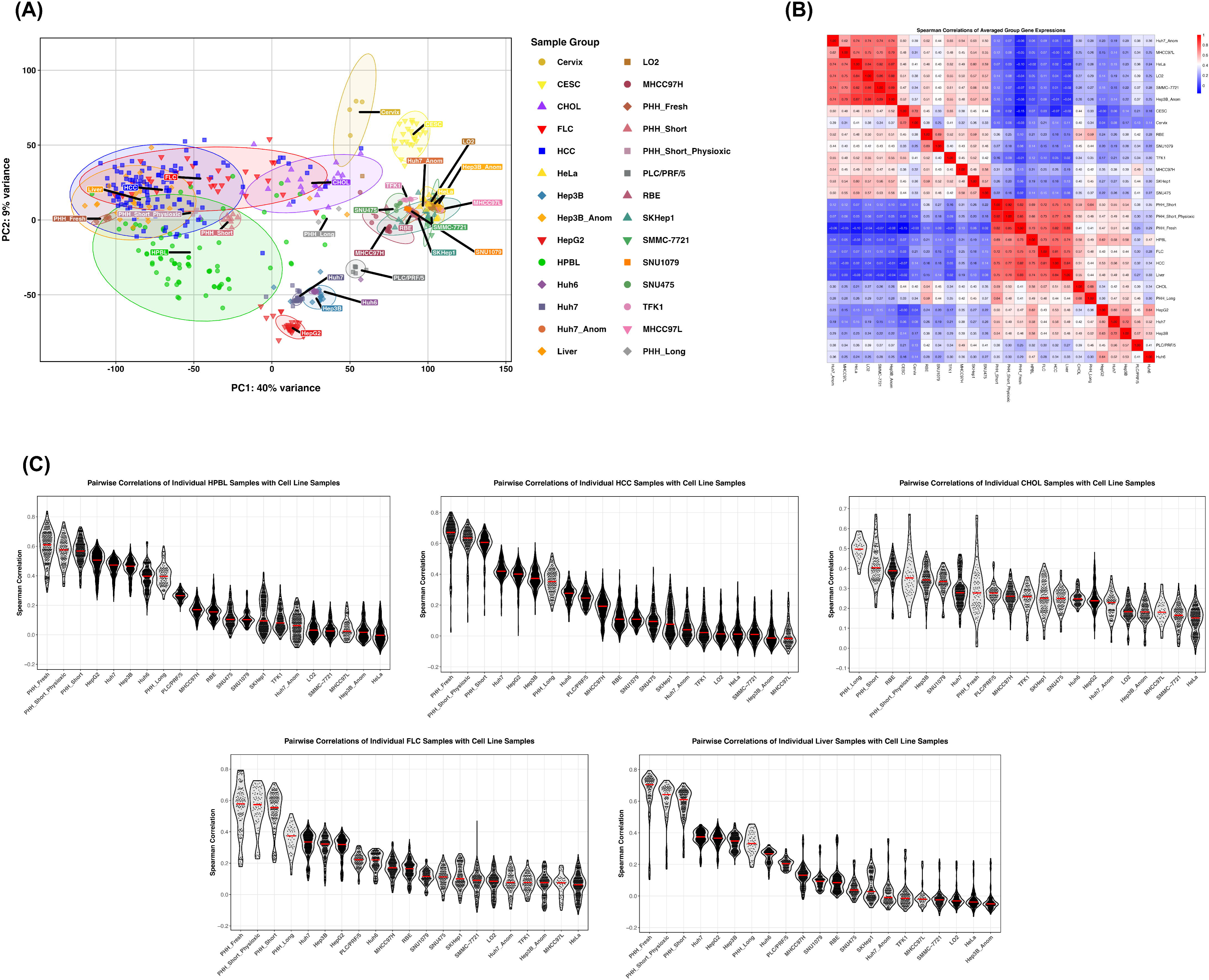
Global transcriptional relationships between liver cancer cell lines and primary tumors. (A) Principal component analysis of all samples using the marker gene set, with transparent probability ellipses delineating sample clusters, showing separation between cell lines and primary tissues. (B) Correlation matrix of averaged expression profiles between cell lines and tumor types. (C) Distribution of pairwise Spearman correlations between individual samples for each cell line-tumor type pair, ranked by median correlation. The clear separation between primary tissues and cell lines in transcriptional space reflects fundamental differences imposed by cell culture conditions, while cell line clustering patterns largely validate their reported origins despite notable exceptions in the MHCC97L, LO2, and SMMC-7721 lines.

**Fig 4.**
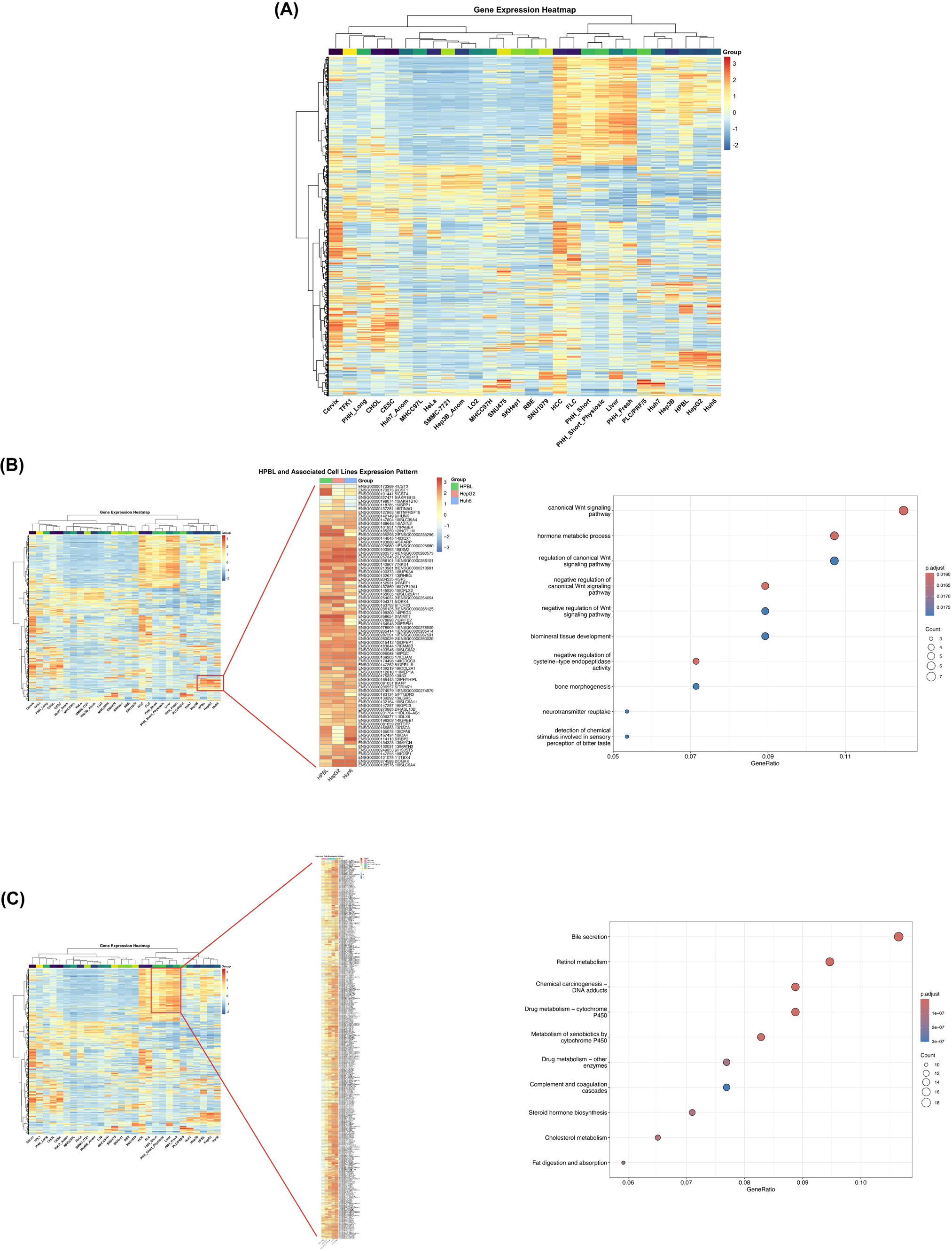
Hierarchical clustering reveals disease-specific expression patterns. (A) Unsupervised hierarchical clustering of all samples using marker genes. (B) HPBL-specific gene expression window showing distinct Wnt pathway activation in HPBL tumors and related cell lines. (C) Progressive loss of liver-specific gene expression in cultured primary human hepatocytes, with partial preservation under physioxic conditions. These distinct expression patterns confirm the HPBL origin of HepG2/Huh6 cells through shared Wnt pathway activation signatures, while highlighting the critical importance of culture duration and oxygen conditions in maintaining hepatocyte-specific transcriptional programs in PHH models.

The PCA revealed a diagonal separation between cell lines and primary tissues across principal components, with cell lines clustering toward the bottom-right and primary tissues toward the upper-left, indicating substantial transcriptional divergence between in vitro and in vivo states.

The correlation matrix quantified these differences through distinct blocks of high inter-correlation within tissue groups and within cell line groups, but lower correlations between these categories. These transcriptional differences likely reflect known culture effects including immortalization, selection pressures, growth on rigid substrates, serial passaging, ambient oxygen exposure, and absence of cellular heterogeneity.^41–43^ Model systems incorporating three-dimensional matrices, co-culture approaches, and physioxic conditions can better recapitulate in vivo characteristics when needed. ^44–46^ While primary liver tissues and cancers formed distinct yet partially overlapping PCA clusters, cell lines showed tight, discrete clustering patterns, reflecting their stable and uniform transcriptional states.

### MHCC97L, LO2, SMMC-7721 and anomalous publicly-available sequencing files demonstrate HeLa-like transcriptomic features

Cervical primary tissues occupied a distinct area in the upper right quadrant of the PCA plot. LO2, MHCC97L, and SMMC-7721 cell lines showed high correlation (0.74 to 0.84) with HeLa cells and clustered within the local HeLa cell PCA region (Figs 3A and 3B), consistent with previous reports of HeLa contamination or derivation.^19^ Notably, the previously identified anomalous Hep3B and Huh7 samples (labeled as Hep3B_anom and Huh7_anom) showed strong correlation (0.74 to 0.88) and regional PCA clustering with the HeLa/LO2/MHCC97L/SMMC-7721 samples (Figs 3A and 3B), as well as inclusion in a broad HeLa-based heatmap cluster (Fig 4A) rather than their purported liver cancer cell line groups, providing additional evidence for their potential misidentification and highlighting the importance of rigorous cell line authentication.

### HepG2 gene expression signatures corroborate HPBL origin

A key finding from our analyses was the strong correlation between HepG2 cells and HPBL samples (Figs 3B, 3C and 4A). This result supports the growing body of evidence asserting that HepG2, often mislabeled as an HCC model, is derived from and more closely resembles HPBL in its gene expression profile.^17^ Both HepG2 and Huh6 clustered with HPBL samples in the unsupervised analyses (Fig 4A), further reinforcing this relationship. While Huh6’s median correlation of all pairwise sample comparisons with HPBL was lower than expected (Fig 3C), its maximum values matched those of HepG2, suggesting potential sample-specific variation. Huh6 cells also displayed a larger gap in correlation with HPBL (0.47) than with HCC (0.34) (Fig 3B).

Closer inspection of the gene expression patterns from the unsupervised clustering analysis revealed a distinctive window of upregulated genes that uniquely differentiate HPBL from other liver cancers, which was also prominently observed in HepG2 and Huh6 cell lines (Fig 4B). Gene Ontology analysis of this HPBL-specific window demonstrated significant enrichment of Wnt signaling pathway components (Fig 4B) encompassing both canonical and regulatory aspects of Wnt signaling. Key genes in this window include AXIN2, DKK4, NOTUM, and SP5, which are all known components or targets of the Wnt signaling pathway, as well as IGSF1, suggesting a complex interplay between Wnt signaling and other developmental pathways in HPBL.^47,48^ Dysregulation of the Wnt signaling pathway through mutations in CTNNB1, which encodes β-catenin, is a well-established driver of HPBL tumorigenesis in patients.^49^ The shared Wnt pathway activation pattern observed between primary HPBL tumors and HepG2/Huh6 cells reflects this clinical reality and aligns with the documented CTNNB1 mutations in both cell lines; HepG2 carries a large in-frame deletion (p.W25_I140del) spanning 116 codons within exons 3 and 4 of CTNNB1, while Huh6 harbors a CTNNB1 point mutation (p.G34V).^49^ These mutations prevent the normal degradation of β-catenin protein, leading to its accumulation and constitutive activation of Wnt target genes. The preservation of this characteristic HPBL molecular signature in HepG2 and Huh6 cells provides transcriptome-level validation of their HPBL origin and supports their utility as relevant in vitro models for studying HPBL biology.

### FLC and CHOL Display Distinct Transcriptional Features with Implications for Cell Line Selection

FLC samples occupied an intermediate position between HCC and CHOL clusters in the PCA space (Fig 3A). This transcriptional "bridging" potentially reflects the complex histological features of FLC, which exhibits both hepatocellular and neuroendocrine characteristics.^50^ This positioning might provide molecular support for a hypothesis that FLC’s unique features and rarity arise from combinatory transformation events affecting multiple unique cell lineages or progenitor cells with mixed differentiation potential.^50,51^ The transcriptional profile of FLC suggests a complex etiology potentially involving simultaneous or coordinated transformation of both hepatocellular and biliary/endothelial lineages, which may explain its distinctive clinicopathological characteristics. For FLC correlations, after the expected ranking of PHHs, the cell lines Huh7, Hep3B, and HepG2 followed a similar pattern to other liver cancers (Figs 3B and 3C). However, the generally lower absolute correlation values with these cell lines highlight a critical gap in FLC research—the lack of well-established immortalized cell line models for this cancer type.

Interestingly, CHOL samples exhibited closer transcriptional similarity to CESC than other hepatocellular-based tissues did, likely reflecting shared epithelial characteristics and transcriptional programs between cholangiocytes and cervical epithelia. This observation was mirrored in the cell lines, where reported CHOL-derived cell lines (TFK1, RBE, and SNU1079) showed increased correlation with HeLa cells and CESC compared to other liver cancer cell lines, but while maintaining sufficient transcriptional distinctness to still form their own unique clusters. The correlation matrix revealed particularly high similarity between RBE and SNU1079 cells (correlation coefficient 0.69), supporting their overlapping clustering in the PCA. SK-Hep-1 cells formed a wider cluster overlapping with CHOL-derived cell lines, with moderate correlation coefficients (0.52 to 0.59) to these groups, reinforcing previous studies that identified these cells as liver sinusoidal endothelial in origin rather than a model for HCCs.^52^

This pattern may present opportunities to leverage existing knowledge from cervical cancer research in understanding CHOL biology, particularly regarding epithelial-specific processes and targeted therapeutic approaches.^53^_–55_ However, it is important to note that CHOL samples still maintained stronger absolute correlation with other liver cancers, suggesting preservation of organ-specific transcriptional programs.

CHOL samples additionally revealed an interesting pattern where long-term cultured PHHs ranked first in correlation, followed by the cholangiocarcinoma-derived RBE cells and comparable correlations with SNU1079 and Hep3B (Figs 3B and 3C). The high correlation with long-term cultured PHHs, especially in comparison to HPBL and HCC correlations with long-term cultured PHHs, is particularly intriguing in light of recent evidence that some intrahepatic cholangiocarcinomas (ICCs) can arise from mature hepatocytes through trans-differentiation. This process has been shown to be Notch2-dependent in mouse models, where hepatocyte-specific deletion of Notch2 can switch ICC-like tumors to a hepatocellular phenotype.^56^ The transcriptional similarities we observe between CHOL samples and long-term cultured PHHs might reflect this biological plasticity, potentially capturing aspects of the hepatocyte-to-cholangiocyte transition that can occur during CHOL development and EMT influences in long-term PHH cultures. Among cell lines, while RBE and SNU1079 were derived from cholangiocarcinomas, the comparable correlation of Hep3B could also reflect this biological phenomenon of cellular plasticity in liver cancer (Fig 3).

### Transcriptional Analysis Reveals Optimal HCC Cell Line Models and Potential Metastatic Signatures

Out of cell lines, HCC samples showed the strongest correlations with Huh7, followed by HepG2 and Hep3B (Figs 3B and 3C). Hep3B displayed notably wider correlation ranges across HCC samples, similar to the patterns observed between Huh6 and HPBL samples where median was lower than expected but maximum was in-line with expectations, suggesting potential subtype-specific modeling considerations. Taking into account the unsupervised clustering analysis, Huh7, Hep3B, and PLC/PRF/5 formed a distinct HCC-associated cluster (Fig 4A), supporting their HCC origin. However, a deeper examination of PLC/PRF/5 cells revealed interesting divergences from the other HCC cell lines in this cluster. While PLC/PRF/5 grouped with Huh7 and Hep3B in broad clustering analyses, they showed relatively lower correlations to primary HCC samples (Figs 3B and 3C) and instead displayed higher correlation patterns with MHCC97H, SNU475, and SK-Hep1 (Fig 3B). These correlation patterns may reflect PLC/PRF/5’s known aggressive phenotype and metastatic potential, as demonstrated by their ability to form rapidly growing tumors with pulmonary metastases in nude mice.^57^

This metastasis-associated transcriptional pattern provides an interesting context for our observations regarding MHCC97H and SNU475 cells. While these reportedly HCC-derived lines cluster together in unsupervised analyses (Fig 4A), they show surprisingly low correlation with primary HCC samples and established HCC cell lines (Figs 3A-C). For MHCC97H, this discrepancy may be explained by its derivation through selective pressure for metastatic potential in mice, suggesting it might better represent metastatic disease than primary HCC.^58^ This observation, combined with their transcriptional divergence from hepatocyte-like states, raises important considerations for their experimental utility. While potentially valuable for studying metastasis, they may not optimally represent primary HCC biology. An alternative explanation for their transcriptional profile comes from possible trans-differentiation of mature hepatocytes to CHOL-like phenotypes as previously mentioned ^56^, which may relate to the clustering of MHCC97H and SNU475 one branch away from CHOL cell lines RBE and SNU1079. It’s crucial to reiterate that the related MHCC97L line shows possible evidence of HeLa contamination in our analyses and should be considered distinct from MHCC97H.

### Physioxic and short-term culture conditions retain similarity to native hepatic state in PHHs

Primary human hepatocytes (PHHs) correlated most closely with normal liver tissue, validating their use for studying normal liver function and metabolism (Figs 3B, C and 4A). However, PHH transcriptional fidelity varied substantially with culture conditions and duration. Fresh PHHs exhibited the highest correlation with primary liver samples (0.83), while short-term cultured PHHs maintained strong correlation (0.75-0.76), suggesting preservation of liver-specific characteristics. PHHs cultured under physioxic conditions (7% O_₂_) demonstrated higher correlation and clustering with liver tissue compared to those cultured under standard atmospheric conditions (21% O_₂_), consistent with previous reports of enhanced preservation of liver-specific markers including albumin and cytochrome P450 enzymes under lower oxygen conditions.^59^ In contrast, long-term cultured PHHs showed markedly decreased correlation with primary liver samples (0.44), approaching levels of liver cancer cell lines. This rightward PCA shift and correlation decline highlights the critical impact of culture duration on maintenance of liver-specific characteristics, suggesting progressive de-differentiation due to adaptation to the in vitro microenvironment.

This pattern was particularly evident in a distinct gene set showing high expression in liver tissue and fresh PHHs but progressive decline during prolonged culture, which was partially mitigated by physioxic conditions (Fig 4C). KEGG pathway analysis revealed significant enrichment (p.adj < 1e-07) in liver-specific metabolic and functional pathways, including bile secretion (GeneRatio = 0.105), retinol metabolism (0.095), chemical carcinogenesis-DNA adducts (0.09), cytochrome P450-mediated drug metabolism (0.09), metabolism by other enzymes (0.08), complement and coagulation cascades (0.08), steroid hormone biosynthesis (0.075), cholesterol metabolism (0.065), and fat digestion and absorption (0.06). Individual gene analysis revealed coordinated downregulation of liver-specific factors, including metabolic enzymes (CYP2C19, CYP8B1, CYP7A1), transport proteins (SLC10A1, SLCO1B1), and glucose homeostasis factors (G6PC1, PCK1). The systematic loss of these genes suggests coordinated dedifferentiation rather than random transcriptional drift, implying that physioxic culture conditions better preserve hepatic gene expression and cellular identity than standard atmospheric oxygen conditions.

Overall, the combined use of PCA, correlation matrix analysis, pairwise Spearman correlations, and unsupervised hierarchical clustering provides a comprehensive view of the relationships between primary tissues and cell lines. By integrating these analytical approaches, we can better understand the suitability of various cell lines as models for specific liver cancers and normal liver function, inform experimental design, and potentially identify novel aspects of tumor biology such as cellular plasticity and trans-differentiation processes from a transcriptomic perspective. The analysis also emphasizes how cell line derivation methods and selective pressures can significantly impact transcriptional programs, potentially moving them away from their tissue of origin.

### PLAGE Analysis Reveals Differential Preservation of Functions Across Cell Lines

To further characterize the relative functional traits between cell lines and primary tissues, we performed Pathway Level Analysis of Gene Expression (PLAGE) using several curated gene sets representing key cancer-related and liver-specific functions (Fig 5). This analysis provided quantitative insights into how well different cell models recapitulate specific aspects of liver biology. Analysis of the HSIAO_LIVER_SPECIFIC_GENES signature (Fig 5A) revealed a clear gradient of liver-specific gene expression across sample types. Fresh primary human hepatocytes (PHH_Fresh) and normal liver tissue showed the highest PLAGE scores, followed closely by PHHs cultured under physioxic conditions. This pattern validates the maintenance of liver-specific transcriptional programs in freshly isolated hepatocytes and further supports the notion that physiological oxygen levels help preserve hepatic character in culture conditions as seen earlier. Most liver cancer cell lines showed substantially lower PLAGE scores, with HepG2 and Huh6 cells demonstrating intermediate scores relative to other cell lines. As expected, HeLa and HeLa-derived lines (including Hep3B_Anom and Huh7_Anom variants) showed the lowest expression of liver-specific genes.

**Figure 5.**
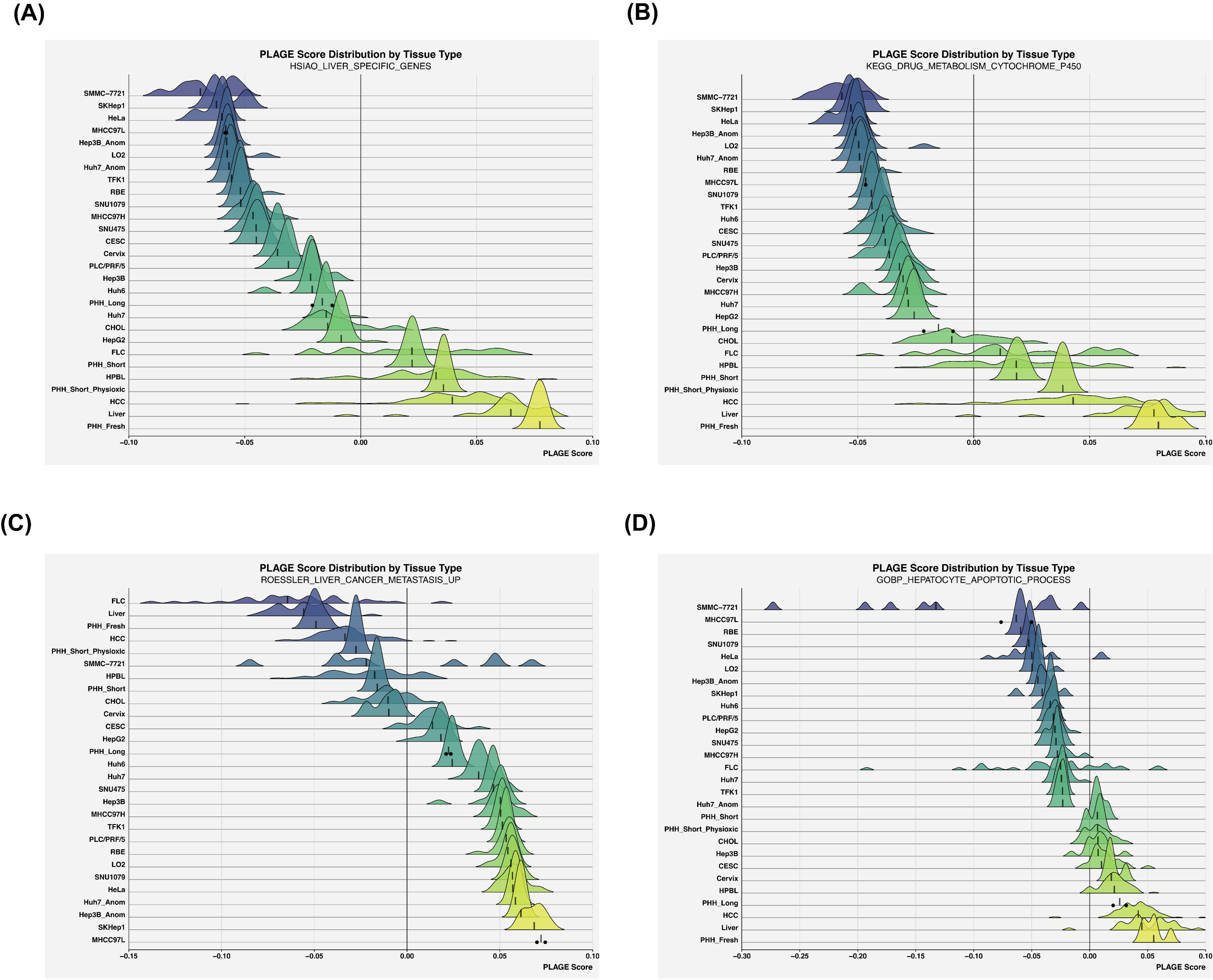
PLAGE analysis reveals functional preservation across cell lines and tumor types. Pathway Level Analysis of Gene Expression (PLAGE) scores showing relative pathway activities. (A) HSIAO_LIVER_SPECIFIC_GENES signature showing gradient of hepatic program maintenance. (B) KEGG_DRUG_METABOLISM_CYTOCHROME_P450 pathway activity demonstrating metabolic competency differences. (C) ROESSLER_LIVER_CANCER_METASTASIS_UP signature revealing enhanced metastatic features in certain cell lines. (D) GOBP_HEPATOCYTE_APOPTOTIC_PROCESS pathway highlighting distinct regulation in cholangiocarcinoma lineages. These PLAGE analyses reveal that while immortalized liver cancer cell lines lose substantial metabolic capacity compared to primary hepatocytes, HepG2 and Huh7 retain intermediate drug metabolism function suitable for initial screening applications, and cell lines with elevated metastasis signatures (PLC/PRF/5, MHCC97H, SNU475) may better model aggressive disease phenotypes.

The KEGG_DRUG_METABOLISM_CYTOCHROME_P450 gene set analysis (Fig 5B) revealed similar stratification but with more pronounced differences between primary tissues and cell models. Fresh PHHs and liver tissue showed markedly higher PLAGE scores than all other samples, highlighting the challenge of maintaining xenobiotic metabolism capacity in culture.

Among immortalized cell lines, HepG2 and Huh7 cells maintained relatively higher PLAGE scores compared to other cancer cell lines, aligning with their common use in drug metabolism studies and toxicity screening.^10,11^ This intermediate metabolic capacity, while reduced compared to primary hepatocytes, makes these lines valuable tools for initial high-throughput drug screening applications in the context of cancer phenotype modeling. However, while most cell lines showed overall negative PLAGE scores, HPBL and HCC tissues maintained intermediate positive scores, suggesting partial retention of metabolic function in tumor tissue that is more extensively compromised in derived cell lines. These findings support a tiered approach to drug metabolism studies, where HepG2 or Huh7 cells might serve as an initial screening tool, followed by validation in more metabolically competent systems such as short-term/physioxic PHHs or stable CYP-expressing cell lines.

Analysis of the ROESSLER_LIVER_CANCER_METASTASIS_UP signature, representing genes associated with liver cancer progression, showed an inverse pattern (Fig 5C). Cell lines, particularly those of HCC origin, demonstrated higher PLAGE scores compared to primary tissues. Among HCC cell lines, PLC/PRF/5, MHCC97H, and SNU475 exhibited notably elevated scores compared to other HCC-origin lines, consistent with their distinct correlation patterns observed earlier and suggesting they may represent more metastasis-like models due to selection and establishment pressures. Notably, PHHs and normal liver tissue showed the lowest scores, consistent with their non-transformed state. Interestingly, FLC samples, while showing considerable variability, displayed the lowest metastasis signature scores among cancer types. This observation appears to align with the relatively low rate of distant metastases in FLC, though it presents an apparent paradox given the high rates of lymph node metastases reported in clinical studies.^60^ This discrepancy may reflect an important sampling bias - many of these case and histological studies rely on surgically resected specimens. While HCC patients with lymph node metastases are typically deemed inoperable, surgical resection remains a standard approach for FLC patients even with extrahepatic involvement.^61^–63 Thus, our metastasis signature findings may better reflect the molecular features of resectable tumors rather than the full spectrum of disease presentation. Alternatively, ROESSLER_LIVER_CANCER_METASTASIS_UP gene signature may be more reflective of distant rather than local metastatic potential.

The GOBP_HEPATOCYTE_APOPTOTIC_PROCESS gene set revealed distinct clustering of cell types based on their apoptotic pathway expression (Fig 5D). Primary liver tissue and fresh PHHs showed elevated PLAGE scores, while most cell lines demonstrated negative scores, potentially reflecting selection for apoptosis resistance in culture. Interestingly, cholangiocarcinoma (CHOL) samples and their derived cell lines (RBE, SNU1079) trended separately from other liver cancers, suggesting distinct regulation of apoptotic programs in this tumor type.

### Assessing Heterogeneity Within Cancer Types and Matching Cell Lines

We finally investigated molecular subtypes within each liver cancer type and assessed how well the highest-matching cell line models recapitulate these distinct tumor phenotypes. For each cancer type, unsupervised clustering was performed using highly variable genes identified through variance modeling across individual samples. Cluster-specific gene signatures were characterized through GO enrichment, KEGG pathway analysis, and GSEA to identify enriched biological processes, altered signaling networks, and differentially activated hallmark pathways. Cell line modeling capacity was evaluated by calculating correlation coefficients between each cell line’s expression centroid and tumor cluster centroids, visualized through spider plot analysis. This subtyping analysis revealed distinct molecular classes within each liver cancer type, with varying representation by established cell lines (Figs 6-9).

**Figure 6.**
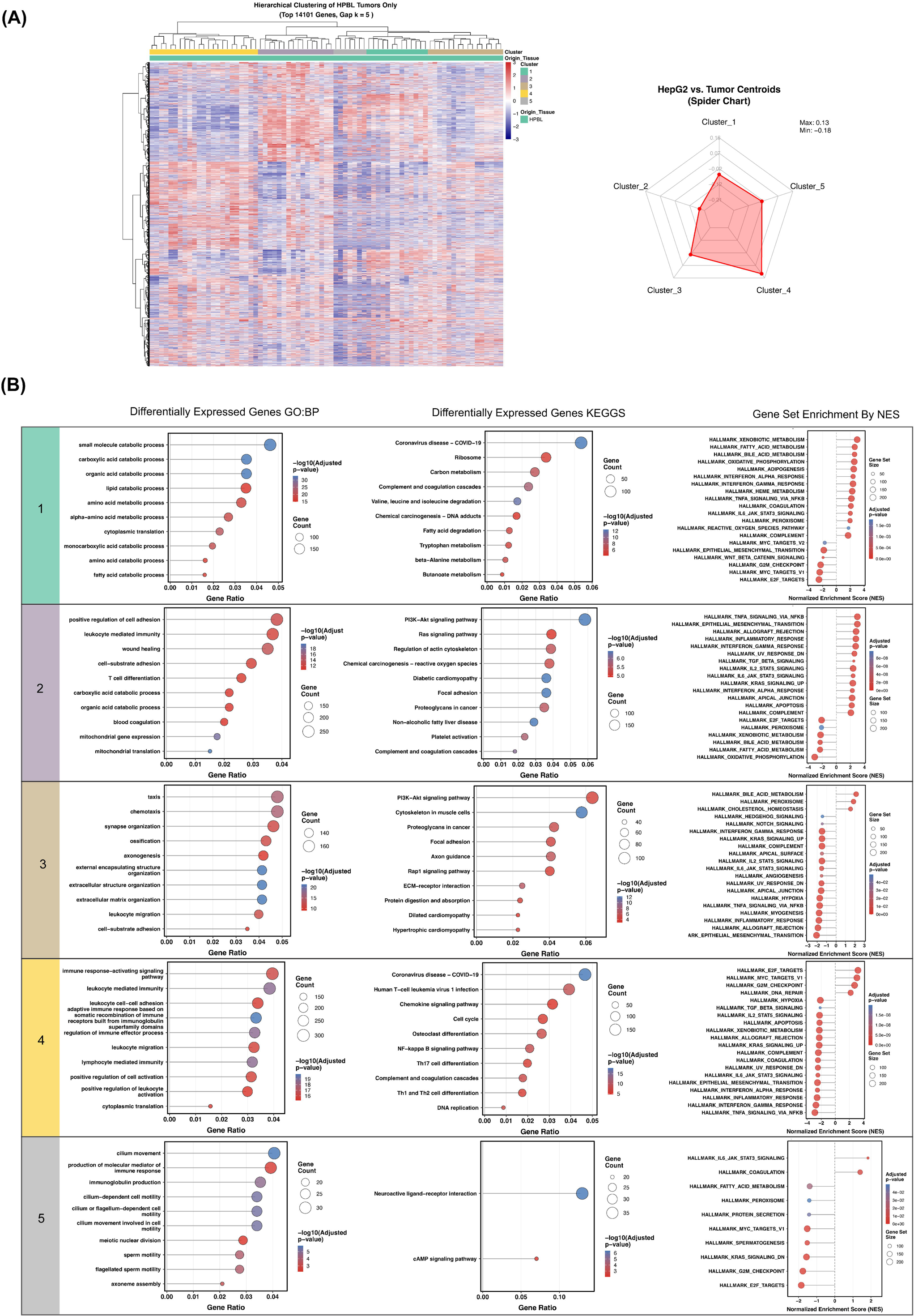
Molecular subtype analysis of hepatoblastoma reveals four distinct clusters. (A) Unsupervised clustering heatmap of HPBL samples based on top variable genes (left) and spider plot showing correlation of HepG2 and Huh6 cell lines with identified HPBL subtypes (right). (B) GO, KEGGS, and GSEA characteristics of each HPBL cluster: Cluster 1 - preserved hepatic function, Cluster 2 - inflammatory signaling, Cluster 3 - extracellular matrix remodeling, Cluster 4 - proliferative immune-suppressive phenotype, and Cluster 5 - altered developmental signaling. (C) Pathway enrichment analysis showing distinct biological processes in each subtype. HepG2 cells show strongest correlation with the proliferative, immune-suppressive Cluster 4 HPBL subtype characterized by high E2F/MYC activity, indicating they best model relatively rapidly dividing hepatoblastomas with immune evasion features rather than inflammatory or ECM-remodeling subtypes.

### HPBL Subtypes Show Distinct Proliferative and Immune Signatures with HepG2 Best Modeling Proliferative Features

Unsupervised clustering identified five HPBL molecular subtypes (Fig 6). Cluster 1 exhibited preserved hepatic metabolism (xenobiotic metabolism, fatty acid metabolism, oxidative phosphorylation) with reduced MYC/Wnt signaling. The reduced activation of oncogenic programs like MYC and Wnt/β-catenin, combined with maintained metabolic functions, suggests these tumors may be less dedifferentiated compared to other subtypes. This interpretation aligns with previous studies showing that hepatoblastoma subtypes exist along a spectrum of differentiation states, with some tumors maintaining substantial hepatocyte-like features.^64,65^ KEGG pathway analysis supported this view, showing enrichment of normal liver metabolic processes including branched-chain amino acid metabolism and fatty acid catabolism, functions that are often lost or dysregulated during malignant transformation.^66,67^ Cluster 2 showed inflammatory, immune-modulatory signatures with PI3K-Akt and TNFα-NFκB activation, suggesting this subtype may represent HPBL tumors with significant immune cell infiltration or inflammatory signaling.^68,69^ Cluster 3 displayed ECM organization and chemotaxis signaling with suppressed EMT/hypoxia pathways. This profile indicates a subtype potentially characterized by altered tumor-stromal interactions rather than aggressive invasive behavior.

Cluster 4 demonstrated high proliferation (E2F targets, MYC) coupled with immune suppression. This pattern suggests a rapidly proliferating HPBL subtype that may evade immune surveillance.^64,70^ Cluster 5 showed unexpected ciliary/hormonal signaling enrichment with IL-6/JAK-STAT3 activation. This unique profile suggests potential dysregulation of developmental signaling pathways in this HPBL subtype.^71,72^

Spider plot analysis revealed that HepG2 cells showed the strongest correlation with Cluster 4, aligning with its proliferative, immune-suppressive phenotype (Fig 6A). This correlation is consistent with HepG2’s well-documented high proliferation rate and altered cell cycle regulation.^73,74^ The shared enrichment of E2F targets and MYC-driven pathways between HepG2 cells and Cluster 4 HPBL tumors suggests that HepG2 cells may be particularly suitable for studying the mechanisms driving rapid cell division and immune evasion in this HPBL subtype. However, researchers should note that HepG2 cells may not optimally represent the inflammatory or ECM-related features characteristic of other HPBL subtypes.

### HCC Subtypes Reveal Distinct Metabolic and Immune States

Unsupervised clustering of HCC samples identified five distinct molecular subtypes (Fig 7). Cluster 1 exhibited an immune-activated phenotype with enriched immune response pathways, platelet activation, and osteoclast differentiation (KEGG), alongside proliferation signatures including E2F targets, MYC, and mTORC1 activation (GSEA). Cluster 2 showed dysregulated lipid metabolism with enriched cholesterol and steroid biosynthesis, altered fatty acid and amino acid metabolism, and inflammatory signatures including TNFα-NFκB, EMT, and interferon response pathways. Cluster 3 displayed immune-modulatory features with leukocyte adhesion and cytokine signaling enrichment (KEGG), and positive enrichment of MYC targets and oxidative stress responses paired with suppressed inflammatory signatures, suggesting immune evasion mechanisms (GO).^75^ Cluster 4 relatively retained hepatocyte-like characteristics with preserved bile acid metabolism, oxidative phosphorylation, xenobiotic processing, and upregulated complement cascades and ECM remodeling. Cluster 5 demonstrated extensive metabolic dysregulation, particularly in catabolic and fatty acid pathways with altered bile secretion and peroxisome function, coupled with pro-inflammatory IL6-JAK-STAT3 and NFκB signaling and suppressed hepatic functions, indicating dedifferentiation.

**Figure 7.**
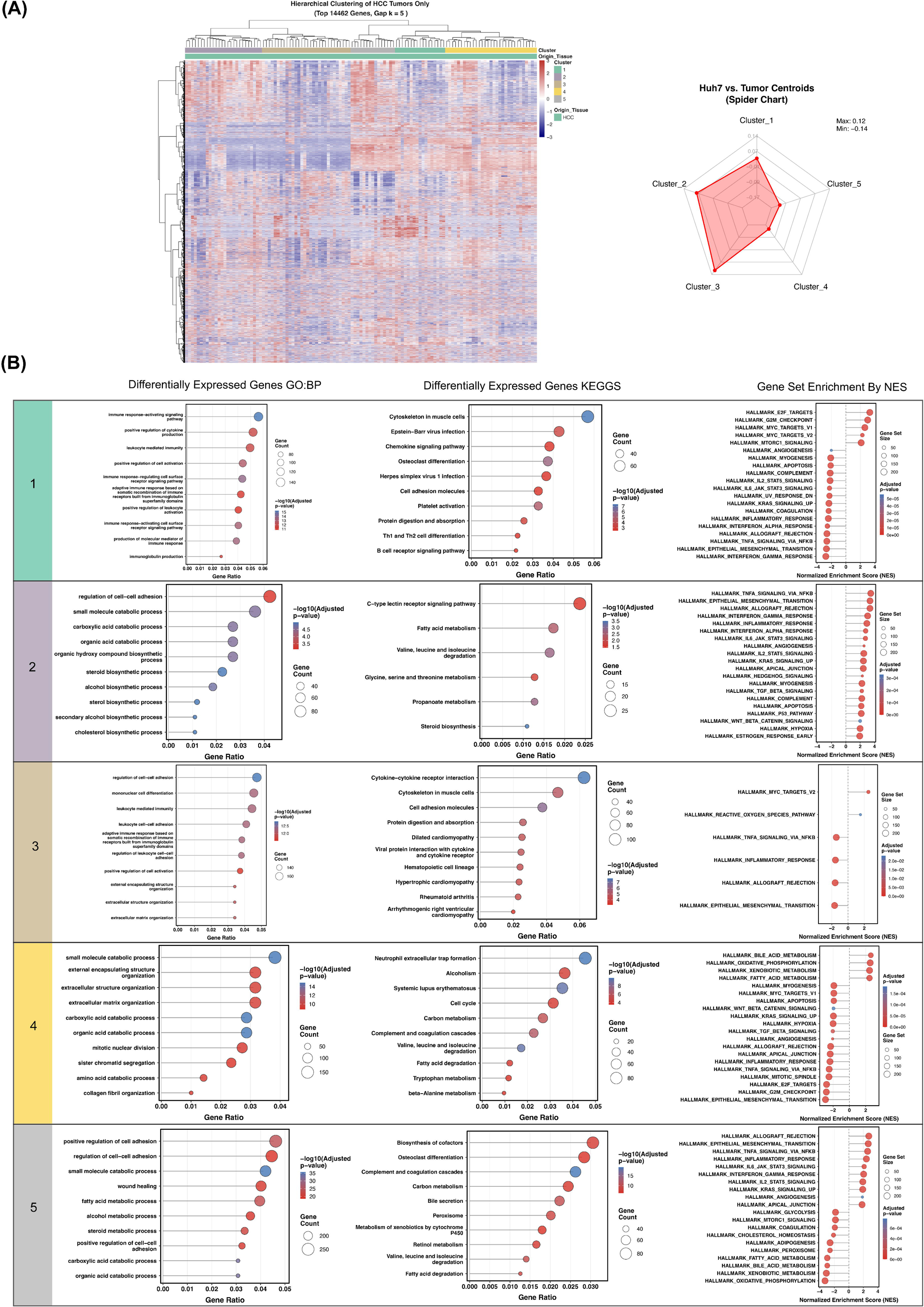
Molecular subtype analysis of hepatocellular carcinoma reveals five distinct clusters. (A) Unsupervised clustering heatmap of HCC samples based on top variable genes (left) and spider plot showing correlation of Huh7 cells with identified HCC subtypes (right). (B) GO, KEGG, and GSEA characteristics of each HCC cluster: Cluster 1 - immune-activated phenotype with enhanced proliferation, Cluster 2 - dysregulated lipid metabolism and inflammatory signaling, Cluster 3 - immune-modulatory with MYC activation, Cluster 4 - preserved hepatic functions and ECM remodeling, and Cluster 5 - metabolic dysregulation with pro-inflammatory signaling. Huh7 cells best model the MYC-activated, immune-evasive HCC subtype (Cluster 3) with impaired inflammatory responses, but show minimal correlation with the more differentiated, metabolically-intact Cluster 4 HCC tumors.

Correlation analysis between Huh7 cells and these HCC subtypes revealed strongest similarity to Cluster 3 (Fig 7A), suggesting they may be particularly suitable for studying Myc-related and immune-modulatory aspects of HCC. Previous studies report impaired TLR3 signaling and high PD-L1 expression contributing towards Huh7 dampening of immune responses.^76,77^ Conversely, Huh7 showed minimal correlation with Cluster 4, indicating departure from the relatively more well-differentiated, metabolically-typical HCC tumor phenotypes.

### FLC Subtypes Display Diverse Metabolic and Neuroendocrine Features with Limited Cell Line Models

FLC clustering identified five distinct subtypes (Fig 8). Cluster 1 showed loss of hepatic metabolic functions with enriched steroid, alcohol, and cholesterol metabolism (GO), bile secretion, cytochrome P450 metabolism, and retinol metabolism (KEGG). GSEA revealed elevated ER stress, unfolded protein response, interferon-alpha signaling, angiogenesis, MYC targets, and PI3K-Akt-mTOR activation, with suppressed NF-κB signaling, suggesting a proliferative, inflammatory, dedifferentiated subtype with invasive potential. Cluster 2 exhibited mitochondrial oxidative phosphorylation enrichment (GO, KEGG) with increased bile acid and xenobiotic metabolism but decreased glycolysis (GSEA), indicating retention of normal liver metabolism without heavy Warburg effect reliance. Downregulated MYC targets suggest reduced proliferative capacity, potentially representing a less transformed phenotype. Cluster 3 demonstrated loss of metabolic plasticity with enriched organic acid catabolism, fatty acid oxidation, and peroxisome function (GO, KEGG), but suppressed xenobiotic metabolism, IL-2/STAT5, interferon-alpha, EMT, MYC targets, and DNA repair (GSEA), suggesting a slow-growing or dormant state with immune evasion capabilities. Cluster 4 showed unexpected neuroactive pathway enrichment—olfaction, sensory perception, neuroactive ligand-receptor interaction (GO, KEGG)—with mildly suppressed pancreatic beta-cell function (GSEA). Given prior reports of neuroendocrine differentiation in FLC tumors ^50^, as well as overlap with bile duct markers^51^, this cluster may represent a neuroendocrine-like subset within FLC tumors. Cluster 5 exhibited stemness and DNA repair features with enriched neuron differentiation and spliceosome regulation (GO, KEGG), plus upregulated MYC targets, DNA repair pathways, and unfolded protein response (GSEA), suggesting genomic stability mechanisms potentially conferring therapeutic resistance.

**Figure 8.**
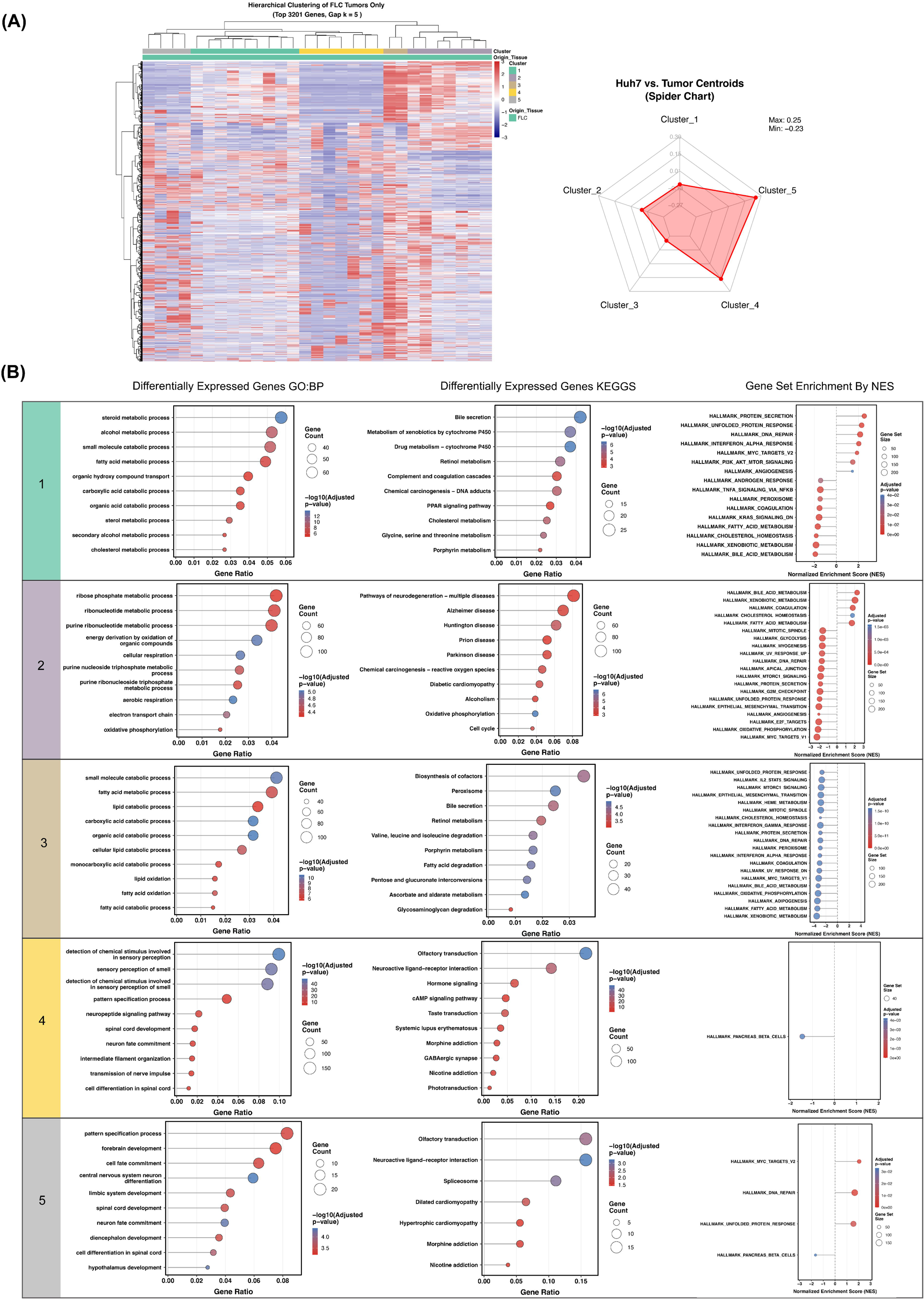
Molecular subtype analysis of fibrolamellar carcinoma reveals five distinct clusters. (A) Unsupervised clustering heatmap of FLC samples based on top variable genes (left) and spider plot showing correlation of Huh7 cells with identified FLC subtypes (right). (B) GO, KEGG, and GSEA characteristics of each FLC cluster: Cluster 1 - metabolic reprogramming with ER stress, Cluster 2 - oxidative phosphorylation and preserved metabolism, Cluster 3 - suppressed immune activation and metabolic plasticity, Cluster 4 - neuroendocrine-like differentiation, and Cluster 5 - stemness features with DNA repair activation. Overall, FLC tumors display remarkable molecular heterogeneity spanning from metabolically active to neuroendocrine-like phenotypes, with the closest matching cell line from a different origin, Huh7 cells, best modeling the proliferative, DNA repair-activated subtype (Cluster 5) but poorly representing the metabolically quiescent subtypes, highlighting the need for additional FLC-specific cell line models.

**Figure 9.**
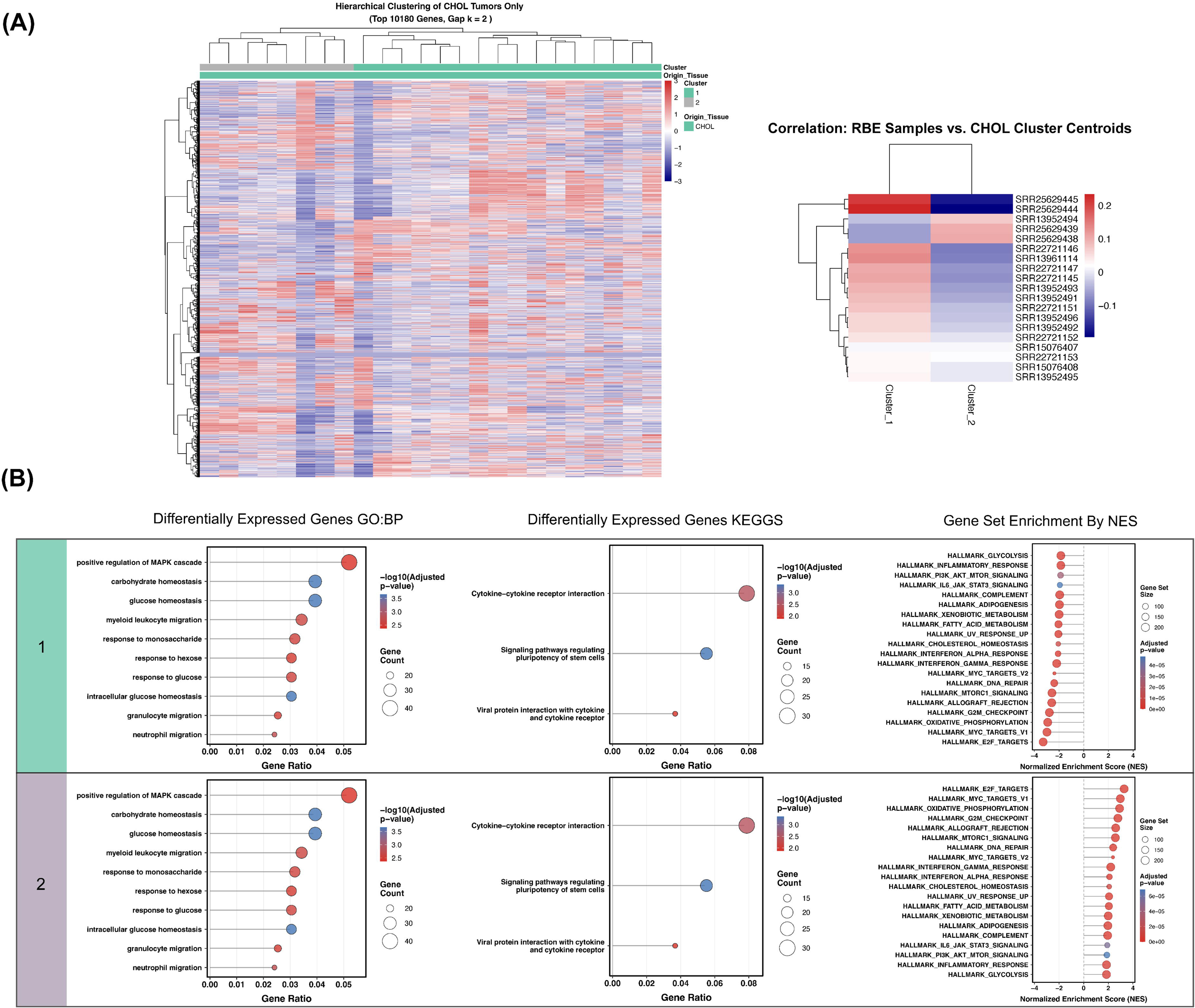
Molecular subtype analysis of cholangiocarcinoma reveals two distinct clusters. (A) Unsupervised clustering heatmap of CHOL samples based on top variable genes (left) and spider plot showing correlation of RBE cells with identified CHOL subtypes (right). (B) GO, KEGG, and GSEA characteristics of each CHOL cluster: Cluster 1 - loss of differentiation with immune evasion, and Cluster 2 - preserved differentiation with higher metabolic and proliferative activity.

Comparison of Huh7 (not derived from a FLC, but closest matching cell line in this analysis scope) transcriptomic profiles with the FLC tumor clusters revealed the strongest correlation with Cluster 5 and the weakest correlation with Cluster 3 (Fig 7A). This suggests that Huh7 cells best model the proliferative, Myc-related, and DNA repair-activated subtype of FLC tumors, whereas they do not as optimally recapitulate the metabolic quiescence and immune-evasive features of Cluster 3.

### CHOL Comprises Two Major Subtypes with RBE Cells Modeling Immune-Evasive and De-differentiated Features

Finally, CHOL analysis revealed no highly-variable multiple clustering patterns based on the selected samples in our set, but they were able to be broadly categorized into 2 major clusters (Fig 9). Cluster 1 is characterized by loss of normal differentiation programs, metabolic suppression, and immune evasion. GO enrichment highlighted significant dysregulation in glucose homeostasis and carbohydrate metabolism, while GSEA revealed relative suppression of hallmark glycolysis and oxidative phosphorylation pathways. Notably, key proliferative pathways, including PI3K-AKT-mTOR signaling and IL6-JAK-STAT3 signaling were also downregulated, consistent with a low-turnover phenotype. KEGG analysis suggested that while stem cell-related signaling pathways were detected, these were likely disrupted rather than actively maintained. The downregulation of inflammatory response pathways and complement signaling further supports a role for immune evasion in this cluster.

CHOL cluster 2, in contrast, retains transcriptional programs associated with normal differentiation and higher metabolic and proliferative activity. This cluster exhibited significant enrichment in MYC and E2F target genes, alongside activation of oxidative phosphorylation and G2M checkpoint regulation. The presence of allograft rejection-related pathways suggests a more immune-permissive state, further distinguishing it from the immune-suppressed profile of Cluster 1. Together, these findings suggest that Cluster 2 may represent a more differentiated, transcriptionally stable subtype, whereas Cluster 1 is characterized by a loss of normal functional programs, metabolic rewiring, and immune escape.

Importantly, RBE cells, which showed the highest correlation with CHOL overall, aligned most closely with Cluster 1, suggesting that this cell line models the more transformed, immune-evasive subtype of CHOL tumors.

## Discussion

Our study presents a comprehensive transcriptomic analysis framework to evaluate the molecular similarities between liver cancer cell lines and their primary tumor counterparts. This approach, while focused on the RNA level, offers unique advantages for investigating cellular phenotypes and disease modeling. The transcriptome-centric methodology provides particular strength in assessing non-coding RNA species, including lncRNAs, and direct transcriptional regulation, which often escape detection in protein-based analyses.^78^ The integration of our findings with existing knowledge of liver cancer subtypes and cell line characteristics provides a robust foundation for model selection. However, we acknowledge the inherent limitations of a transcriptomic-only approach. Post-transcriptional modifications and protein-level regulation cannot be directly assessed through RNA sequencing alone, although advancements in some technologies may help bridge these gaps.^79^ Additionally, bulk RNA-seq data at steady-state provides a snapshot rather than temporal dynamics of gene expression and the dynamic range of cellular responses that might be relevant for specific experimental contexts. As such, this analysis is intended to be used with other sources of information as part of a comprehensive framework for selecting appropriate cell line models, incorporating factors such as mutation profiles, epigenetic states, metabolic characteristics, and functional assays specific to the research question at hand.

Our transcriptomic analysis revealed critical insights regarding cell line authentication, PHH culture conditions, and model system selection. Initial quality control identified potential authentication concerns with specific Hep3B and Huh7 samples displaying correlation patterns characteristic of HeLa cells rather than their purported liver cancer origins. These anomalous samples clustered with known HeLa-derived cell lines in correlation analyses and principal component projections, suggesting contamination or misidentification. Such findings reinforce the ongoing challenges of cell line authentication in cancer research ^20^ and highlight how transcriptome-wide analyses can serve as a complementary validation approach alongside standard authentication methods. The implications extend beyond individual studies: misidentified samples in public repositories, annotated as HCC cell lines, risk propagating incorrect conclusions through secondary analyses, meta-analyses, and machine learning approaches. These findings emphasize rigorous sample validation in both primary research and curation of public genomic databases.^20,25^

PHH analysis revealed distinct time and oxygen-dependent dynamics in gene expression. Fresh PHHs exhibited highest correlation with primary liver tissue (r = 0.83), followed by short-term cultured PHHs (r = 0.75-0.76). PHHs cultured under physioxic conditions (7% O_₂_) demonstrated better preservation of liver-specific transcriptional programs than standard atmospheric conditions, consistent with previous studies.^59^ Long-term cultured PHHs showed marked correlation decrease (r = 0.44), approaching liver cancer cell line levels. Progressive loss of liver-specific characteristics in genes involved in bile acid synthesis, xenobiotic metabolism, and glucose homeostasis suggests coordinated dedifferentiation rather than random transcriptional drift, emphasizing the importance of culture duration in experimental design.

HepG2 analysis provided strong transcriptional evidence supporting HPBL origin while revealing value for specific HCC research aspects. HepG2 cells showed strongest correlation with HPBL samples (r = 0.62) and clustered consistently with HPBL tissue, aligning with recent studies identifying hepatoblastoma origin.^17^ However, high ranking across both HPBL and HCC correlations may reflect retention of core hepatocyte-like features. This dual nature presents both challenges and opportunities: while use as a general HCC model requires caution, retained hepatocyte-like characteristics suggest previous HepG2 studies generated valuable insights into liver biology, even if erroneously interpreted in an HCC context rather than HPBL. This understanding allows for more informed experimental design, where HepG2 cells can be employed strategically based on their actual biological properties, and previous research findings can be appropriately recontextualized within the framework of HPBL biology while still maintaining relevance to broader hepatic processes.

PLAGE analysis offers a framework for model selection based on research objectives. While HepG2 cells correlate strongly with HPBL in Wnt signaling pathways, they maintain sufficient liver-specific metabolic gene expression to model certain HCC biology aspects, highlighting the importance of considering both pathway-specific similarities and broader transcriptional patterns. Molecular subtyping analyses further refine this framework by revealing distinct disease clusters with unique pathway activation patterns. HepG2 cells best model the proliferative, immune-suppressive HPBL subtype with enhanced Wnt signaling but are less suitable for inflammatory or ECM-focused HPBL subtypes. Huh7 cells show strongest correlation with the immune-modulatory, MYC-activated HCC subtype, indicating optimal use for studying these specific HCC aspects.

While this study provides insights into transcriptional similarities between liver cancer cell lines and primary tumors, several important clinical considerations were outside our analytical scope. Our analysis did not account for potential confounding variables in the primary tumor samples such as patient age, ethnicity, and sex, which can influence tumor biology and gene expression patterns. These demographic factors could impact the transcriptional profiles of primary tumors and, by extension, their correlation with cell line models. Future analyses should consider these variables in their experimental design and analysis to provide more precise context-dependent recommendations for cell line selection.

## Conclusions and Future Directions

Based on our comprehensive transcriptomic analysis of liver cancer cell lines and primary tumors, several critical areas require attention to advance the field. One pressing research need is the development of dedicated FLC cell lines as a part of ongoing efforts.^3,81^ Our analysis highlighted a significant gap in FLC modeling, as existing common cell lines showed limited correlation with FLC transcriptional profiles. The unique molecular features of FLC, including its distinctive position between HCC and CHOL profiles, emphasize the importance of developing representative and reproducible in vitro models, which would be invaluable for studying FLC-specific biology and therapeutic responses. Additionally, expanded FLC sample collection, sequencing, and contribution to public repositories would strengthen our understanding of tumor heterogeneity and improve model selection.

Existing model systems require systematic validation using contemporary approaches. While our analysis focused on transcriptional profiles, comprehensive validation should incorporate multiple layers of characterization, including genomic, epigenomic, and metabolomic analyses. This is particularly important for widely used cell lines like HepG2, where our findings reinforced its HPBL origin rather than HCC. Emerging opportunities include the application of machine learning approaches to cell line selection and validation. Recent developments, such as CASCAM ^82^, offer promising frameworks for predicting cell line-tumor matches based on multi-omic data integration. These computational approaches could help researchers select optimal models for specific research questions. These could also scale as part of authentication protocols that require updating to incorporate transcriptional profiling alongside traditional methods. Our identification of potential HeLa contamination in certain samples highlights the limitations of current authentication approaches. Regular authentication using multiple methodologies or integrated pipelines would enhance research reliability both at the primary source of use, and on publicly-available data used for secondary analysis.

Culture conditions significantly impact primary cell phenotype and functionality, as demonstrated by our PHH analysis. Standardization of culture conditions, particularly oxygen levels, passage numbers, and media composition, is crucial for reproducibility. Our observation that physioxic conditions better preserved liver-specific gene expression patterns suggests that standard culture practices may warrant revision. In parallel, novel primary cell culture methods show promise for better maintaining in vivo characteristics. Three-dimensional culture systems, co-culture approaches, and microfluidic devices could better recapitulate native cell state and the tumor microenvironment.^44–46^

When comparing transcriptomic profiles of tumors, tissues, and cell types, researchers should be aware of the following study and experimental design issues that may arise from these samples and this data Key points of consideration include: (1) the specific research question and required cellular characteristics, (2) the transcriptional similarity between cell line and tumor subtype, (3) the stability of relevant pathways under standard culture conditions, and (4) the presence of specific mutations or alterations relevant to the study. Experimental design should incorporate appropriate controls and validation steps; this includes: (1) authentication at periodic intervals, (2) monitoring of passage numbers and phenotypic stability, (3) validation of key findings in multiple cell lines when possible, and (4) confirmation of critical results in primary tissue samples or animal models. Data sharing and reproducibility requirements should include: (1) regular submission of RNA-seq data to public repositories, and (2) authentication protocols on one’s own cell lines in primary research, and on files obtained for secondary analyses and use cases. The implementation of these recommendations would enhance the reliability and translational relevance of liver cancer research. As new technologies and methods emerge, these guidelines should evolve to incorporate improved approaches for model system characterization and validation.

## Supporting information

Supplemental Gene Signature Analysis

Supplemental HPBL Window

Supplemental PHH Window

Supplemental GEO Metadata

## Data Availability Statement

The raw sequencing data analyzed in this study are publicly available through the NCBI Sequence Read Archive (SRA) database (https://www.ncbi.nlm.nih.gov/sra). SRA accession numbers and BioProject identifiers for all samples are provided in the Supplementary Materials. Databases, gene set collections, and annotation resources were accessed as detailed in the Materials and Methods section. Processed data, analysis results, and metadata generated in this study are included in the Supplementary Materials.

## Notes

## Abbreviations

ATCC: American Type Culture Collection
CESC: cervical squamous cell carcinoma
CHOL: cholangiocarcinoma
CV²: coefficient of variation squared
DESeq2: differential expression analysis for sequence count data
ECM: extracellular matrix
EMT: epithelial-mesenchymal transition
ER: endoplasmic reticulum
FDR: false discovery rate
FLC: fibrolamellar carcinoma
GEO: Gene Expression Omnibus
GO: Gene Ontology
GSEA: Gene Set Enrichment Analysis
HCC: hepatocellular carcinoma
HPBL: hepatoblastoma
ICC: intrahepatic cholangiocarcinoma
KEGG: Kyoto Encyclopedia of Genes and Genomes
PCA: principal component analysis
PHH: primary human hepatocytes
PLAGE: Pathway Level Analysis of Gene Expression
RNA-seq: RNA sequencing
STR: short tandem repeat
VST: variance stabilizing transformation.

